# Pairwise interactions and serial bottlenecks help explain species composition in a multi-species microbial community

**DOI:** 10.1101/2024.11.22.624877

**Authors:** Kasturi Lele, Benjamin E. Wolfe, Lawrence H. Uricchio

## Abstract

Characterizing the processes that drive microbial community assembly remains a key challenge in ecology. Several recent studies have argued that pairwise interactions may be insufficient to explain co-occurrence patterns in complex microbial communities, but most such studies have focused on synthetic communities not found in nature or microbes grown in contexts that differ widely from their natural environment. Moreover, most models of pairwise interactions rely on equilibrium assumptions that are not relevant to all natural communities, such as gut microbiomes or species in fluctuating environments. Inclusion of appropriate demographic factors into models of pairwise interactions could be a potential approach to better capture patterns of community assembly. In this study, we investigated whether multi-species co-occurrence patterns can be predicted from pairwise interactions for microbes isolated from sourdough starters. Interaction parameters inferred from pairwise growth trajectories were suggestive of widespread coexistence between pairs of microbes in our species’ pool. In communities of up to nine species, most species’ presence and relative abundance could be reliably predicted based on a model of pairwise interactions. The inclusion of nonequilibrium demography in our model further improved the accuracy of our pairwise model. Our work contributes to the broader debate on the processes underlying community assembly by showing that pairwise interactions are predictive of community structure in a system of moderate species complexity.

## Introduction

Species interactions are major determinants of ecological communities and characterizing the processes that underlie community assembly remains a major goal of ecology research. Several (non-exclusive) processes could help explain co-occurrence patterns, including pairwise competition (Friedman, Higgins, and Gore 2017; Venturelli et al. 2018), emergent coexistence (Chang et al. 2023), resource-mediated interactions (Brochet et al. 2021; Tannock et al. 2012; Pozo et al. 2016), and higher-order interactions (Mayfield and Stouffer 2017; Gibbs, Levin, and Levine 2022; Ishizawa et al. 2024). While pairwise interactions seem to be sufficient to explain co-occurrence patterns in some systems (Friedman, Higgins, and Gore 2017; Venturelli et al. 2018; Dedrick et al. 2023), they fail to accurately predict membership in others (Momeni, Xie, and Shou 2017; Chang et al. 2023). Though it remains unclear what drives these disparate findings, potential explanations include difficulties in predicting compositions of more complex communities (May 1972; Allesina and Tang 2012), co-evolutionary history (Wade 2007) or prevalence of mutualistic interactions (Madsen et al. 2016; Foster and Bell 2012). Differences in modeling and parameter inference choices could also play a role.

One major challenge that has limited our ability to characterize the processes driving community assembly is the difficulty of experimentally validating the predictions of theoretical models. Much of the work on mathematical models of community assembly has examined plant and animal communities, where experimental manipulations are possible over short terms but it is difficult to validate findings of models over longer timescales (HilleRisLambers et al. 2012). Microbial communities have substantial roles in determining biodiversity from the population to ecosystem scale, and are more amenable to validation because of the rapid timescale of growth and logistical feasibility of experimental manipulation. Many studies have applied mathematical models to microbial communities (reviewed in (Succurro and Ebenhöh 2018; Kumar et al. 2019; van den Berg et al. 2022). However, even in microbial communities, results supporting (Friedman, Higgins, and Gore 2017; Gralka et al. 2020) and contradicting (Momeni, Xie, and Shou 2017; Sanchez-Gorostiaga et al. 2019) the utility of pairwise interactions as predictors of community membership have been reported. Many prior studies investigated only a subset of culturable microbes from a complex environment or used communities formed by species that may not naturally co-occur (Friedman, Higgins, and Gore 2017; Venturelli et al. 2018). Studies in more naturalistic systems (such as germ-free plants or animals) might better represent naturally occuring patterns of microbial diversity, but are often less straightforward to manipulate experimentally than synthetic communities (Jans and Vereecke 2025; Bosch, Guillemin, and McFall-Ngai 2019; Yi and Li 2012; Seedorf et al. 2014). Additional studies using more realistic and tractable microbial communities are needed to understand when and how species interactions can predict community assembly.

Microbial communities derived from fermented foods are experimentally tractable systems that can provide opportunities to measure and model the causes and consequences of species interactions (Wolfe and Dutton 2015; Wolfe 2018). One of the oldest fermented foods we know of is sourdough, prepared from a mixture of grain flour (usually wheat) and water, and used to make bread (Minervini et al. 2014; Van Kerrebroeck, Maes, and De Vuyst 2017; Calvert et al. 2021; De Vuyst, Comasio, and Van Kerrebroeck 2023). Sourdough starter production and consumption is widespread across the globe, and thus it is important to be able to manipulate and control the growth of its constituent microbial communities. Over the past decade, several studies have cataloged and demonstrated the bewildering diversity of microbes across sourdough starter cultures (De Vuyst and Neysens 2005; Minervini et al. 2014; Oshiro, Zendo, and Nakayama 2021). Though studies have observed a diversity of microbes across different sourdough starter cultures, a single starter usually has between 3-5 bacterial species and 1-2 yeast species. This combination of a small community size and a diverse microbiome sampled across different starter cultures allows for a variety of interactions to occur, while being able to parse the mechanism of interactions and track changes over longer time scales. Thus, we can use this system to ask questions about microbial community assembly for a co-evolved set of species in the context of their natural environment.

Though differences in co-evolutionary history and species complexity may drive the heterogeneous findings on pairwise or higher-order interactions in microbial communities, modeling choices could also have a substantial impact. Most pairwise modeling approaches use equilibrium assumptions to compute expected patterns of community membership, but non-equilibrium demographic processes could also play a role in determining community structure and might help explain differences between pairwise models and observed patterns of diversity (Cenci and Saavedra 2019). Many microbial species experience environmental fluctuations that affect abundance and could play a role in determining community membership. For instance, like most laboratory microbes used in microbial ecology research, sourdough communities are periodically transferred into fresh substrate in kitchens and bakeries (De Vuyst, Comasio, and Van Kerrebroeck 2023). Periodic disturbances can destabilize communities at equilibrium (Philippot Laurent, Griffiths Bryan S., and Langenheder Silke 2021), but also could increase the likelihood of coexistence in some pairwise competitions (Schreiber, Benaïm, and Atchadé 2011; Hening, Nguyen, and Chesson 2021). Thus, the inclusion of realistic demographic factors in pairwise models has the potential to improve their fit to observed patterns without necessarily needing to invoke higher-order interactions.

In our study, we investigated whether a classic ecological model for pairwise interactions (the generalized Lotka Volterra model, or gLV) can explain observed co-occurrence patterns of microbes isolated from sourdough starters. We expected coexistence to be prevalent between microbes in this system, since each of these microbes is adapted to the abiotic environment and many are known to co-occur in sourdough starters (Landis et al. 2021). However, since microbes were isolated from different starter cultures, niche overlap might result in competitive exclusion between some pairs of species. We built a nine-species gLV model and parameterized it using growth rates and interaction coefficients from single and pairwise growth curves for all species in our species’ pool to ask whether observed co-occurrence patterns can be predicted in this natural system. We assembled experimental communities composed of the full set of nine microbial species, as well as communities leaving out one species at a time, to compare with the predictions of our model. Finally, we also tested whether adding demographic factors in the form of repeated bottlenecks would alter the predictions of coexistence in the gLV model.

## Methods

### Experimental methods -

#### Forming the species pool for the experiments -

Members and collaborators of the Wolfe lab previously conducted a survey of several sourdough starters from across the world (Landis et al. 2021). From this collection, they isolated and experimentally characterized four lactic acid bacteria (*Fructilactobacillus sanfranciscensis, Levilactobacillus brevis, Lactiplantibacillus plantarum*, and *Companilactobacillus paralimentarius)* and four yeasts (*Saccharomyces cerevisiae, Wickerhamomyces anomalus, Kazakhstania humilis*, and *Kazakhstania servazzii*) We have used the traditional names for species in the genus *Kazakhstania* rather than recently proposed taxonomic rearrangements in order to preserve continuity with earlier sourdough literature (Liu et al. 2024). These microbes were chosen as they were the most abundant microbes across different sourdough starter cultures, and showed strong positive or negative patterns of co-occurrence among themselves. In this study, we used an additional acetic acid bacterium (*Acetobacter malorum*) from the same collection along with these eight microbes, as the role of these bacteria in the sourdough community has not been studied widely (Rappaport et al. 2024).

#### Measurement of microbial growth over time (growth curves) -

For both single species and pairwise growth curves, we grew all microbes in a liquid cereal-based fermentation medium (CBFM) using a previously developed protocol (Landis et al. 2021). We picked this medium as we wanted to use a liquid medium that would match the composition of the dough environment that the sourdough microbes were isolated from. Briefly, we mixed 100% whole wheat and all-purpose flour in equal proportion by mass, suspended this mixture in deionized water (1 part flour: 9 parts water by mass), and allowed the solids to settle by centrifuging the mixture at 3000 rpm for half an hour. We then collected and filtered the supernatant through Falcon filter funnels (0.20 μm pore size). We acknowledge that this medium doesn’t have the full suite of carbohydrates available in traditionally made sourdoughs, but is nevertheless able to fulfill the growth requirements of our microbes. For the growth curve experiments as well as multi-species community assembly experiments, we standardized the input inocula by diluting 15% glycerol stocks (stored at −80°C) for which the CFU concentration was known with 1X phosphate buffered saline (PBS).

For single species growth curves, we standardized input inocula to 1,000 CFUs per 10 μl for each of the 9 species in the species pool. This inoculum was added to 240 μl of CBFM in individual 1.5 ml microcentrifuge tubes (USA Scientific, FL, USA), with 3 experimental replicates for each growth curve. When the cultures were inoculated, we measured the total species abundance at input (0 hrs). The cultures were incubated at 24°C and allowed to grow statically for a total of 68 hrs with the cap sealed. Within this growth period, we homogenized the cultures and measured their total species abundance at the following timepoints - 6 hrs, 18 hrs, 24 hrs, 30 hrs, 42 hrs, 48 hrs, 54 hrs and 68 hrs. All species abundances were measured by CFU counts as described below. Thus, for each growth curve, we had measures of species abundance at 9 time points. This experiment was repeated a second time with three replicates per species.

For pairwise growth curves, we standardized input inocula to 1000 CFUs per 10 μl for every species. For each pair of species, we then mixed the inocula before adding 10 μl of the mixed inoculum to 240 μl of CBFM in individual 1.5 ml microcentrifuge tubes, with 3 measurement replicates for each pair. The final input inoculum for each species was thus standardized to 500 CFU per 10 μl. In total, this experiment had 36 pairs of species for a fully factorial pairwise design. Total species abundance at input (0 hrs) was measured separately for each species. Subsequently, the cultures were allowed to grow statically and total species abundance was measured together at the same time points as the single species growth curves (Figure 1A).

**Fig 1.**
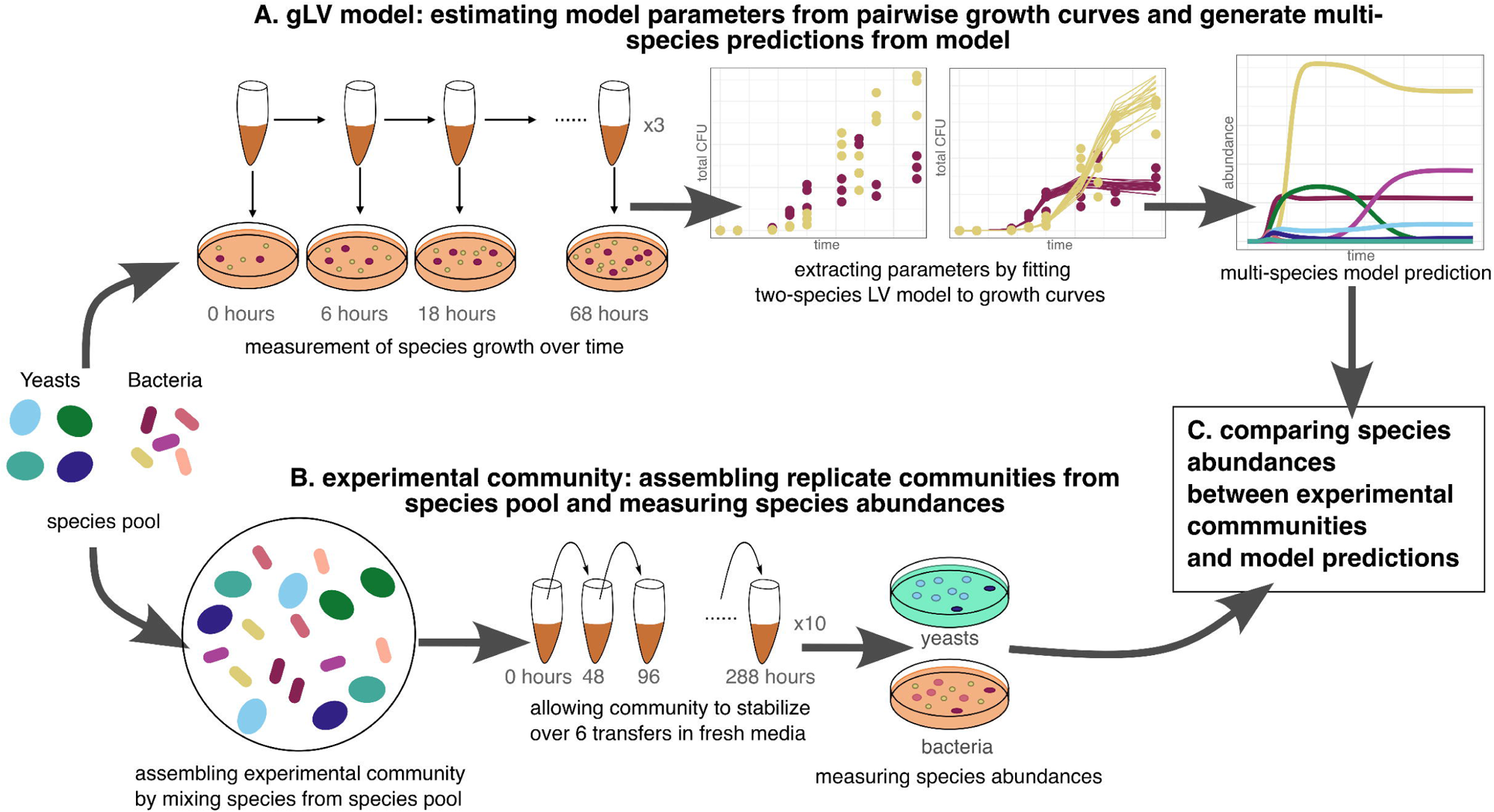
Conceptual figure detailing the experimental procedure. (A) Experimental procedure for obtaining model predictions. We measured growth curves from single and pairwise combinations of all our species in our species’ pool, and obtained growth parameters by fitting two-species gLV models to these growth curves. We then used these parameters to get predictions of species abundance from the 9-species gLV model. We also obtained predictions of pairwise coexistence from the growth parameters. (B) Experimental procedure for the experimental observations. We assembled experimental communities by mixing species from the species pool. We created a 9-species community and nine 8-species communities, leaving out one species in turn. We measured species’ abundance at the end of six transfers, to compare with the gLV model predictions. (C) Comparing species abundances from both gLV model and experimental communities. We used species’ ranks in communities and Bray-Curtis dissimilarities between communities to compare species’ presence and abundance between model and experimental communities.

For each determination of total species abundance, 5 μl of culture was serially diluted up to a 10^-5^ dilution, spot-plated (3 μl of each dilution from 10^-1^ to 10^-5^) on nutrient agar, and colonies were allowed to grow for 2-5 days before counting CFUs. For the single species growth curves, bacteria were grown on MRS agar (Hardy Diagnostics, CA, USA) and yeasts were grown on YPD agar. For the pairwise growth curves, bacteria-bacteria pairs were grown on MRS agar and differentiated using colony morphology. Yeast-yeast pairs were grown on WL nutrient agar (MiliporeSigma, MA, USA) and differentiated using colony color and morphology. Bacteria-yeast pairs were grown on selective media (YPD with added chloramphenicol (50 mg/L) for yeast, and MRS with added natamycin (21.6 mg/L) for bacteria) to quantify CFU. For pairwise growth curves, the abundance of a given species was assigned to be 0 if it fell below the detection limit of 1/100th of the total population at a given time point. For examples of colony morphologies of different bacteria and yeast species, see Appendix S1: Figure S1.

#### Multi-species community assembly experiment -

To test whether multi-species communities would assemble in a similar manner as predicted in the model, we set up experimental communities with all 9 species, as well as leave-one-out communities (LOO) with 8 species, leaving out one species each time from the original species’ pool. For all community assembly experiments, we used CBFM as well as standardized inocula as described above.

For all multi-species communities, we standardized input inocula to 1,000 CFU per 10 μl for each species. 10 μl of this inoculum was added to 190 μl of CBFM in individual 1.5 ml microcentrifuge tubes. 10 experimental replicates were established for each community (9-species as well as all LOO communities), for a total of 100 communities. These communities were incubated at 24°C and allowed to grow without shaking. They were transferred to fresh CBFM every 48 hours, adding 10 μl of culture into 190 μl of CBFM. In this manner, we conducted six transfers over two weeks. At the end of the sixth transfer, we made 15% glycerol stocks for all communities, and stored them at −80°C for further analysis.

To measure abundance of each species at the end of the experiment, 20 μl of the community stored as glycerol stock was serially diluted up to 10^-6^ dilution and plated on selective agar to differentiate bacteria from yeast. We checked the species-specific survival rates post freeze-thaw to make sure that none of the species had particularly low survival after one round of freeze-thaw, similar to the conditions in our experiment. Bacteria were counted on MRS agar plates with added natamycin, and different species were identified using colony color and morphology. Yeasts were counted on WL nutrient agar plates with added chloramphenicol, and different species were identified using colony color and morphology (Figure 1B). For both yeasts and bacteria, we confirmed the identity of species with similar colony morphologies by extracting DNA from putative colonies using the DNEasy Powersoil Pro kit (Qiagen, PA, USA), and amplifying the ITS region (for yeasts) and 16S region (for bacteria) using PCR. We then obtained DNA sequences for the amplified regions using Sanger sequencing (Genewiz from Azenta, MA, USA), and species’ identities were confirmed both with comparisons to reference sequences of previously characterized strains as well as a Blast search in the NCBI database using the Geneious Prime platform (Version 2023.2.1). We confirmed the identity of the following species in this manner - *S. cerevisiae*, *K. humilis*, and *C. paralimentarius*.

### Computational methods -

#### Inference of Lotka-Volterra model parameters -

To extract growth rates and interaction coefficients from the growth curve data, we fit one-or two-species gLV models to the growth curves measured experimentally. We used a slightly modified form of the gLV equations (Mühlbauer et al. 2020), specifically

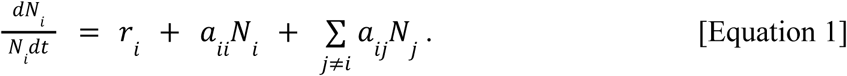

*N_i_* represents the abundance of species *i*, *r_i_* is the basal growth rate, and *a_ii_* and *a_ij_*are the coefficients for intraspecific interaction (interaction of species *i* with itself), and interspecific interaction (interaction of species *i* with all other species *j*). Note that the summation term on the right hand side represents interspecific interactions and vanishes for single species growth curves.

We used a two-step rejection sampling approach to infer model parameters. First, we inferred approximate posterior distributions of *r_i_*and *a_ii_* using single species growth curves; then, we constrained these two parameters by treating their posteriors as priors in the second round of inference and inferred *a_ij_* from pairwise growth curves. Specifically, we sampled *r_i_* and *a_ii_*from their inferred posterior distributions by randomly picking from the values we obtained in the first round of inference. We used this constrained approach because the *r_i_* and *a_ii_* terms appear in both the single species growth equations for species *i,* as well as every pairwise competition experiment that involves species *i.* Any pairwise growth experiment in which species *i* was rapidly outcompeted by species *j* contains little information about species *i*’s parameter values, and using all pairs involving *i* simultaneously for the inference of *r_i_* and *a_ii_* would result in fitting to a very large number of summary statistics simultaneously (Figure 1A).

We wrote custom scripts in Python (3.11.4) to estimate parameters from growth curves. First, we randomly sampled values of *r_i_*,*a_ii_*, and *a_ij_* using the numpy and random libraries. We sampled 5*10^6^ parameter combinations for single-species growth curves and 10^7^ parameter combinations for pairwise growth curves. We picked the initial values of *N_i_* and *N_j_* from the observed growth curves. Then, we obtained the parameters that best fit our observed data by minimizing the log-squared distance between the modeled and observed growth curves at all time points, given by

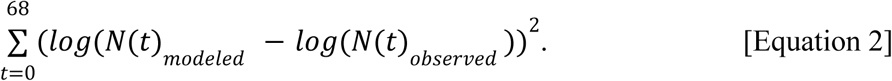

We constrained the values of *a_ij_* to be negative to prevent unbounded growth in multi-species model predictions – biologically, this corresponds to an assumption that interactions between species are primarily competitive rather than facilitative. We acknowledge that this would prevent the detection of cross-feeding among some pairs of microbes. We made this choice to avoid unbounded growth in the multi-species model, and believe that the niche overlap-fitness difference framework used later would still allow us to capture the existence of positive interactions between microbes in our system. These parameters were used to compute pairwise coexistence outcomes and to predict multispecies community composition. The experimentally observed species abundances and the best-fitting growth trajectories we used to extract *r_i_*, *a_ii_*, and *a_ij_* values can be seen in Appendix S1: Figure S2. We used the parameter values from 20 best-fitting trajectories in both steps of parameter inference for the theoretical predictions of coexistence as well as for multi-species model predictions. The distributions of *r_i_*, *a_ii_*, and *a_ij_* values can be seen in Appendix S1: Figure S3.

#### Building a gLV model of multi-species community -

We wrote a script in Python (3.11.4) that used the solve_ivp function from the SciPy library (Virtanen et al. 2020) to numerically solve the full set of time-dependent gLV equations using empirically-inferred parameter values. For a given set of parameters, the script solved the gLV differential equations, producing species abundances for all species as output. We obtained solutions for the set of differential equations at the final time t = 288 hours for all model simulations presented in this paper, which was the final time point at which we sampled species abundances in the multi-species community assembly experiment. For the core set of predictions for the 9-species community, we ran this script over different combinations of parameters to get 100 model predictions (Figure 1A). Additionally, to isolate the impact of pairwise effects, we also ran a version of the script where *a_ij_* values were randomly assigned to species, keeping *r_i_* and *a_ii_*values the same (Appendix S1: Figure S4).

#### Theoretical predictions of outcomes of pairwise competition -

We used a theoretical method to generate predictions of stable coexistence for pairs of species (Chesson 1990; Letten, Ke, and Fukami 2017). Briefly, the mutual invasibility criterion gives a simple test for coexistence (Grainger, Levine, and Gilbert 2019): when *a_ii_* > *a_ji_* and *a_jj_* > *a_ij_*species *i* and *j* are predicted to coexist. This criterion can be reframed in terms of the niche overlap between species (ρ) and their average fitness difference (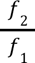) (Letten, Ke, and Fukami 2017). These terms can be written in terms of the interaction coefficients from the gLV model we used as given below. A derivation of how we arrived at these equations from the ones used by Letten et. al. is given in Appendix S1: Section S1.

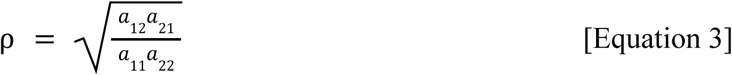

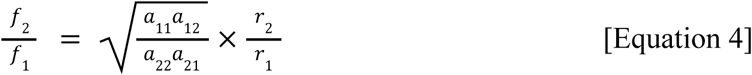

Under the mutual invasibility criterion, species coexist when *a_ii_* > *a_ji_*and *a_jj_* > *a_ij_*, or equivalently

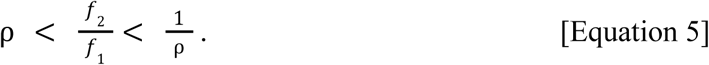

Additionally, if ρ > 1, then alternative stable states are possible for the competing pair, and order of arrival can influence competition outcomes (Ke and Letten 2018). Using this theoretical framework, we calculated values of ρ and 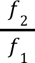 for all pairs of species and made predictions of stable coexistence based on these values. These analyses were performed using a custom R script.

#### Comparison between pairs of species with and without serial bottlenecks -

We investigated how demographic processes, especially serial bottlenecks corresponding to media transfers, might affect community composition in this system. We modeled bottlenecks as periodic reductions in population size, where each species in a metacommunity is sampled in proportion to its current abundance at the sampling time point. To create a version of the model with serial bottlenecks, we made a small modification to the python script we used above. For simulations reported here, we supposed a dilution factor of 1/20 with bottlenecks occurring every 48 hours. This process of dilution and growth is similar to the conditions experienced by our experimental communities. All other aspects of the model code remained unchanged. We used this model in comparison with the older model without bottlenecks with growth parameters for two of the pairs of species in our community, *W. anomalus:C. paralimentarius*, and *A.malorum:L.plantarum*. We sampled 100 parameter combinations from our posterior estimates for each pair. We used the Python module matplotlib.pyplot to plot an informative subset of outcomes from the serial dilution model (Hunter May-June 2007). Additionally, we used this version of the model with serial bottlenecks to obtain model predictions for 8-species communities (leaving out one species in turn from the model) that matched our leave-one-out experiments. For these model predictions, we used the same script but set the initial species abundances to 0 for the excluded species. We conducted similar statistical analyses on these new sets of communities as we did on the earlier 9-species communities.

#### Statistical analyses and figures -

Analyses and plots were done in R (4.3.1 “Beagle Scouts”), using various packages as specified. We used the R packages ggplot (Wickham 2016) and cowplot (Wilke, Wickham, and Wilke 2019) for all our figures, unless stated otherwise. We conducted community analyses using the package vegan (Oksanen 2010)), except where otherwise denoted.

To visualize relative abundances of species for model predicted and experimentally observed communities, we used the function vegan::vegdist to calculate Bray-Curtis dissimilarities, and the stats::hclust function (method = ward.d2) to hierarchically cluster replicate communities. We also ranked species by abundance and calculated the average rank of all species over all replicates for model and experimental communities.

We compared species composition in experimental and model communities by calculating Spearman’s rank correlation on the average rank of model vs experimental species, using the function stats::cor.test (method = ‘spearm’, null hypothesis = true rho is equal to 0). Ties among ranks were resolved by fractional ranking. Species with the same abundance (most commonly, 0 abundance) were assigned a rank that was averaged between the next two ranks. We also compared relative species abundances between model and experimental communities by calculating the Shannon diversity index using the vegan::diversity function. We performed a two-tailed t-test using the function stats::t.test (null hypothesis = difference in means between groups is 0, significance threshold = 0.05) to test for differences in Shannon diversity between these groups. Additionally, we performed a PERMANOVA on Bray-Curtis dissimilarities using the function vegan::adonis2 (permutations = 1000, test statistic = pseudo-F, number of groups = 2, null hypothesis = communities are interchangeable when group identities are permuted) to evaluate what fraction of variation in Bray-Curtis dissimilarities between model predictions and experimental communities are explained by group identity.

To visualize the spread of model vs experimental communities, we conducted NMDS using the function vegan::metaMDS (k = 2, maxit = 999, try = 500). For this analysis, we included 10 replicates for experimental communities and 100 replicates for model communities. From the coordinates obtained in NMDS, we calculated species scores using the function vegan::wascores to approximate the contribution of each species’ to the MDS axes. This function returns the centroid of the species distribution, weighted by species’ abundance in each community. To further understand whether any of the species in our species pool were associated with model or observed communities, we conducted an indicator species analysis using the function indicspecies::multipatt. All these analyses above were repeated for model communities with serial bottlenecks.

For comparisons between community compositions for LOO communities and the 9-species experimental community, we performed a PERMANOVA (function = vegan::adonis2, permutations = 1000, test statistic = pseudo-F, number of groups = 9). We included 10 replicates for the 9-species community and for each of the LOO communities, with the exception of the community without *F. sanfranciscensis*, which had nine replicates. We then used NMDS to visualize the spread of communities using the function vegan::metaMDS (k = 2, maxit = 999, try = 500) and calculated species’ scores using vegan::wascores.

In order to get a broad overview of the model predictions, and compare the overlap between model predictions and experimental communities, we used the distribution of Bray-Curtis dissimilarities. We calculated Bray-Curtis dissimilarities between each model and experimental community. Then, for each model community, we kept only the minimum value of Bray-Curtis dissimilarity from all corresponding experimental communities. We then plotted the distribution of these values to visualize how different model predictions compared with corresponding experimental communities. We also calculated and plotted the distribution of the minimum Bray-Curtis dissimilarity for model predictions with and without serial bottlenecks.

## Results

### A. Coexistence is widespread among pairs of species in the species pool

In order to understand which pairs of species in our species pool were predicted to coexist, we used the niche overlap-fitness difference framework (Letten et. al. 2016) to generate predictions of coexistence, one species outcompeting the other, or the presence of priority effects. We observed that predicted coexistence was prevalent across different pairs (Fig 2A), with 79% of sampled parameter estimates suggesting coexistences. To look for further trends, we subdivided the species pairs into groups by bacteria:bacteria, bacteria:yeast and yeast:yeast. We observed that for bacteria:yeast pairs (cross-kingdom interactions), parameters estimated from growth curves indicated predictions of coexistence slightly more often (84%) than bacteria:bacteria (75%) or yeast:yeast pairs (68%).

**Fig 2.**
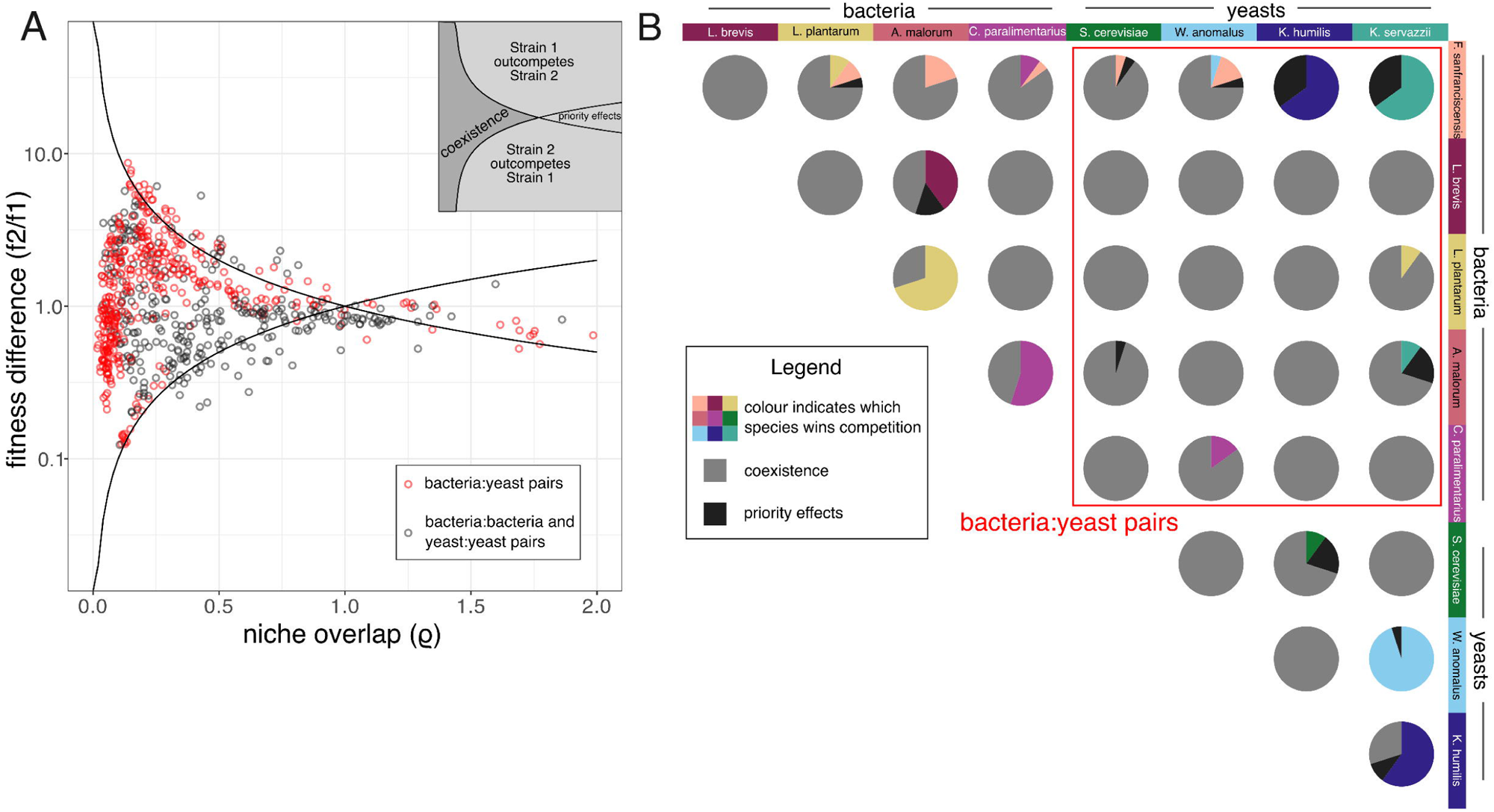
Graphs showing predictions of coexistence across all pairs of species in our species pool. (A) Predictions of coexistence from parameter combinations for all species projected on the niche overlap-fitness difference graph. 712 predictions of coexistence for 36 pairs are shown. Figure inset indicates the region of coexistence (ρ < f_2_/f_1_ < 1/ρ). Red dots indicate predictions of coexistence from bacteria:yeast pairs. (B) Predictions of coexistence split by pairs of species. In each pie chart, predictions of coexistence indicated by light gray, priority effects indicated by dark gray, and the species predicted to win is indicated by its respective color as shown in the column headers. Red box highlights all bacteria-yeast pairs. Overall, most pairs are predicted to coexist, with bacteria:yeast pairs having a slightly higher proportion of coexistence. We then probed deeper into the predictions of coexistence for each pair of species (Fig 2B). For 17 out of 36 species pairs, parameters estimated from different growth curves for the same pair of species gave rise to conflicting predictions of coexistence vs persistence of one species. Interestingly, the remaining 19 pairs were all predicted to coexist with each other for all inferred parameter values, highlighting the widespread coexistence between microbes predicted in this system.

### B. Predictions of community composition from the multi-species gLV model generally align with experimental community compositions

After examining the predictions of widespread coexistence in our system from the theoretical framework, we turned our attention to the predictions from our multi-species generalized Lotka-Volterra (gLV) model. We first looked at the composition of model communities. From the multi-species gLV model, we obtained relative species abundances and hierarchically clustered them using Bray-Curtis dissimilarities to look at the distribution of community composition across 100 iterations of the model (Fig 3A). Across all replicate model communities, we observed that community composition was relatively conserved. In descending order of abundance, model communities consisted of the species *L. plantarum* (average abundance = 55.81%), *C. paralimentarius* (average abundance = 14.27%), *W. anomalus* (average abundance = 11.73%), *L. brevis* (average abundance = 11.63%), *S. cerevisiae* (average abundance = 5.15%), and *K.humilis* (average abundance = 1.42%). When we implemented a detection limit of 1/100 (that was present for experimentally observed communities), the abundance of *F. sanfranciscensis*, *A. malorum* and *K. servazzii* fell below the detection limit for the vast majority of communities (97). Bacteria were far more abundant than yeasts in model communities, making up 81.71% of the community while yeasts constituted 18.29% of the community.

**Fig 3.**
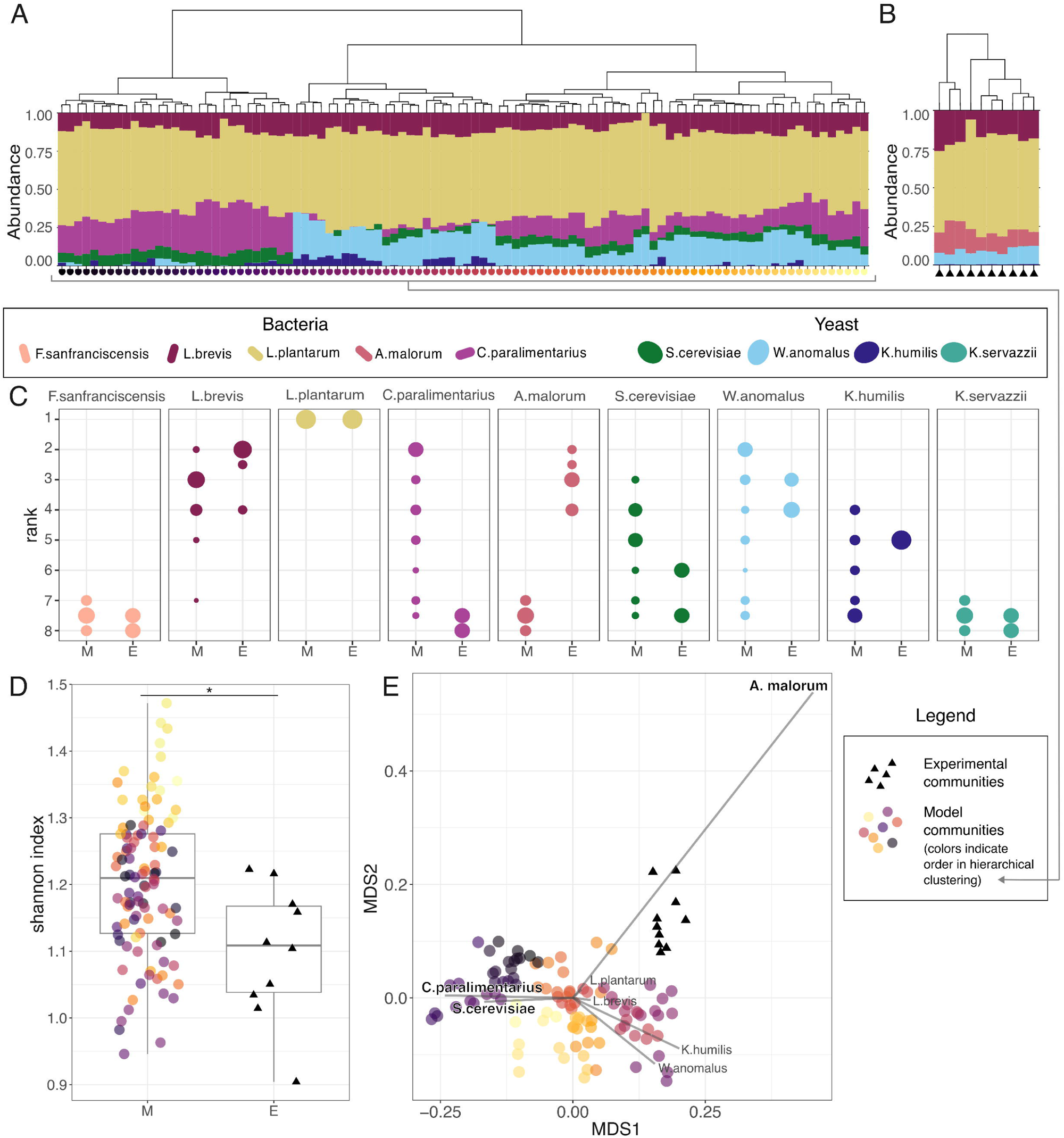
Comparisons of model predictions with experimentally observed communities. (A) Model communities hierarchically clustered by Bray-Curtis dissimilarities. (B) Experimentally observed communities hierarchically clustered by Bray-Curtis dissimilarities. For A and B, height of each bar indicates relative abundance of corresponding species in a given community. Dendrograms above the graphs indicate communities more similar to each other. (C) Distributions of species’ ranks compared between model (M) and experimental (E) communities. Higher ranks indicate greater abundance in the community. Dot size indicates the fraction of communities where species had corresponding ranks. (D) Shannon diversity index between M and E communities. (E) NMDS plot visualizing Bray-Curtis dissimilarities between M and E communities. For D and E, black triangles indicate experimental communities and coloured circles indicate model communities. Colors indicate distance in hierarchical clustering from Fig 3A for model communities. Gray vectors and species names indicate species’ scores.

Next, we investigated how the community composition among experimentally observed communities varied across 10 replicate communities. (Fig 3B). Once again, we observed that community composition was relatively similar across the replicates. In descending order of abundance, experimental communities consisted of the species *L. plantarum* (average abundance = 59.82%), *L. brevis* (average abundance = 17.60%), *A. malorum* (average abundance = 13.21%), *W. anomalus* (average abundance = 8.96%), *K.humilis* (average abundance = 0.38%), and *S.cerevisiae* (average abundance = 0.01%). Overall, bacteria made up 90.36% of the community, while the relative abundance of yeasts was 9.36%, dropping to half of that predicted by the model. Most of this decrease was due to lower numbers of *W. anomalus* and *S. cerevisiae* in the experimental community.

Finally, we compared the community compositions that were predicted by the gLV model and that were observed in the experiment. We observed that overall, most of the same species were present in both model as well as experimental communities. However, relative abundance differed between model predictions and experimental communities. To quantify some of these differences, we calculated the distribution of species’ ranks in models and experiments (Fig 3C). Yeasts were more abundant in model communities, and some bacteria had their ranks flipped (notably, *A. malorum* was absent in model communities, and *C. paralimentarius* was absent in experimental communities). Overall, we found a positive correlation between the average ranks of species in model versus experimental communities, though the correlation was not statistically significant (Spearman’s rho = 0.41, p = 0.27). Using a set of modified model predictions with randomized *a_ij_* values, we observed that species-specific interactions play an important role in determining community structure (Appendix S1: Figure S4).

Despite broad similarities in composition between model and experimental communities, the aforementioned differences suggest that our model predictions may form statistically distinct communities from experimental communities. We used various diversity metrics to quantify these differences. First, we compared the alpha diversity of the two community types by computing the Shannon index (Fig 3D). Despite overall similarities, model communities had significantly higher values of Shannon index, compared to experimental communities (mean = 1.20 for model, mean = 1.10 for experimental, p = 0.0095), indicating that model communities had a more even distribution of species. We also visualized community differences based on the Bray-Curtis dissimilarities between samples using non-metric multidimensional scaling (NMDS) of community structure (Fig 3E), and tested whether these differences between communities were significant using PERMANOVA. Indeed, we found that model and experimental communities cluster separately on an NMDS plot, though the difference between model and experimental communities accounted for 21% of the variation between communities (p < 0.001, partial R^2^ = 0.21). To understand which species were driving these differences in community composition, we calculated weighted average species’ scores and overlaid them on top of the NMDS plot (Fig 3E), observing that *A. malorum* was associated with experimental communities and *C. paralimentarius* was associated with model communities. Additionally, we conducted an indicator species analysis that showed that *S. cerevisiae* and *C. paralimentarius* were indicators for model communities (p=0.001) and *A. malorum* was an indicator for experimental communities (p=0.001). (Appendix S1: Figure S5).

### C. Incorporating repeated bottlenecks in the model may increase prediction efficiencies

We investigated whether demographic processes that may affect the experimental communities could also have an impact on model outcomes. To that end, we added periodic bottlenecks to our model to mimic the transfer into fresh nutrient medium that occurred in our experiment. Bottlenecking reduced the abundance of *C. paralimentarius* and increased *W. anomalus* abundance in a subset of model predictions. In a single combination of parameters sampled from the posterior, *A. malorum* was predicted to be present in the community (Fig. 5A). These changes were reflected in the distribution of Bray-Curtis dissimilarities between the model predictions with and without bottlenecks, where a subset of model predictions now aligned better with experimental observations (Fig. 5C). However, since this change only occurred in a subset of model predictions, we did not observe major changes in the distribution of species’ ranks, or in the NMDS plot of Bray-Curtis dissimilarities for model predictions with bottlenecks. Whereas there was no significant difference between the values of Shannon index between model predictions with bottlenecks and experimentally assembled communities (Fig. S4). Thus, we conclude that serial bottlenecks improved model predictions, but only for a subset of predictions. To understand how repeated bottlenecks might drive these shifts in community outcomes, we examined six illustrative examples pairwise growth with and without repeated bottlenecks from the parameters we obtained from *C. paralimentarius* and *W. anomalus* competitions and *L. plantarum* and *A. malorum* competitions (Fig 6). Note that these combinations of parameters were not sampled randomly, but were selected to represent different possible impacts of serial bottlenecks. We picked these two pairs of species in particular to explore further because model predictions and experimental observations differed for *C. paralimentarius* and *A. malorum* in a subset of model predictions. Additionally, in our theoretical predictions of coexistence, a subset of posterior distribution predicted coexistence while another subset predicted exclusion of *A. malorum* by *C. paralimentarius*. In the subset of plotted parameter combinations, we observe both conserved outcomes (species composition is the same with and without bottlenecks) and divergent outcomes (e.g., competitive exclusion that flips to co-occurrence on the timescale of the experiment for the same set of model parameters). We may observe shifts in community composition in only a subset of model predictions with bottlenecks due to these differences in outcomes for different parameter combinations for the same species pairs.

**Fig 4.**
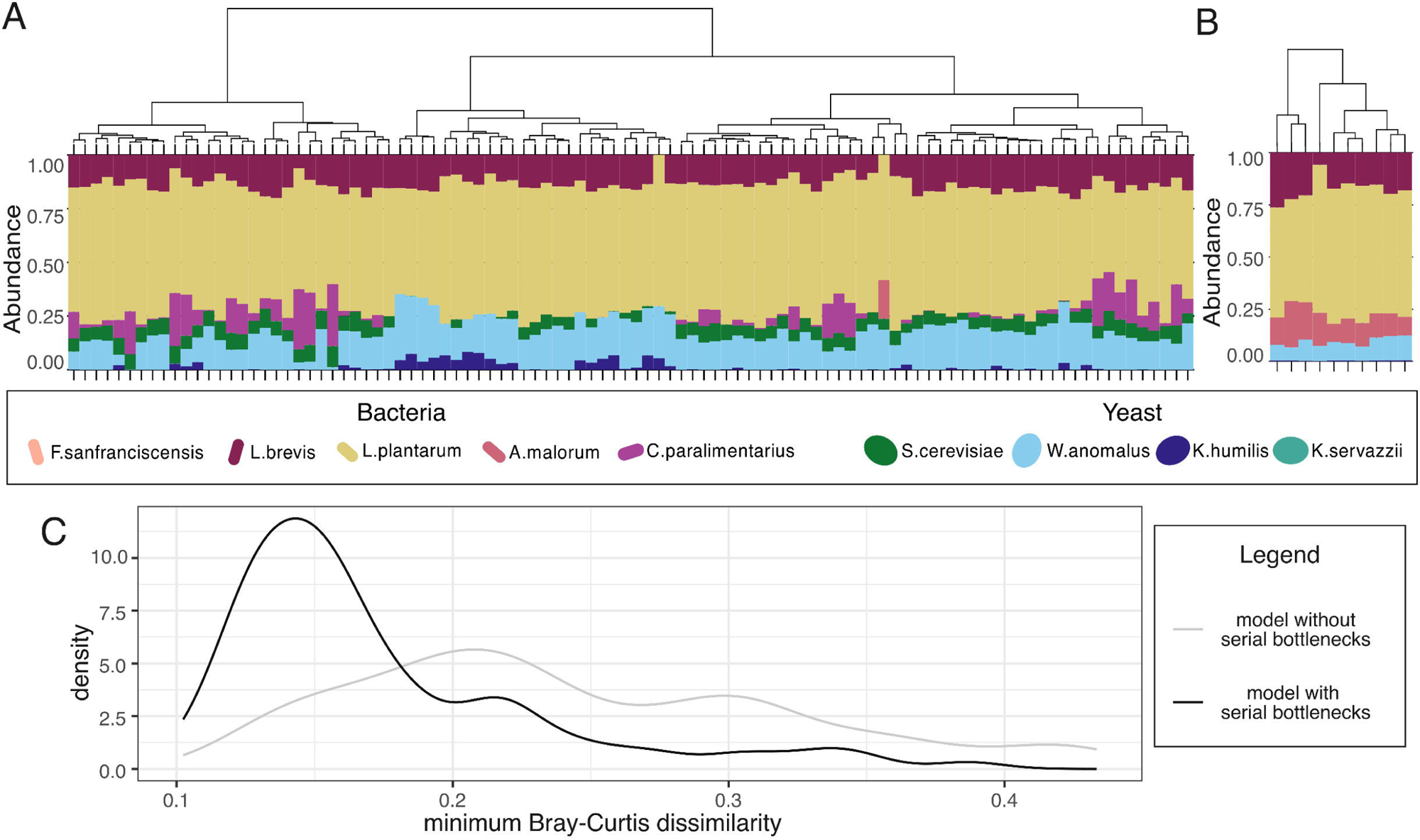
Selected analyses on model predictions with repeated bottlenecks. (A) Model communities hierarchically clustered by Bray-Curtis dissimilarities. (B) Experimentally observed communities hierarchically clustered by Bray-Curtis dissimilarities (replicated here from Fig. 3. for ease of comparison). For A and B, height of each bar indicates relative abundance of corresponding species in a given community. Dendrograms above the graphs indicate communities more similar to each other. C) Density plot showing distribution of Bray-Curtis dissimilarities between all model predictions with (black line) and without (grey line) repeated bottlenecks.

**Fig 5.**
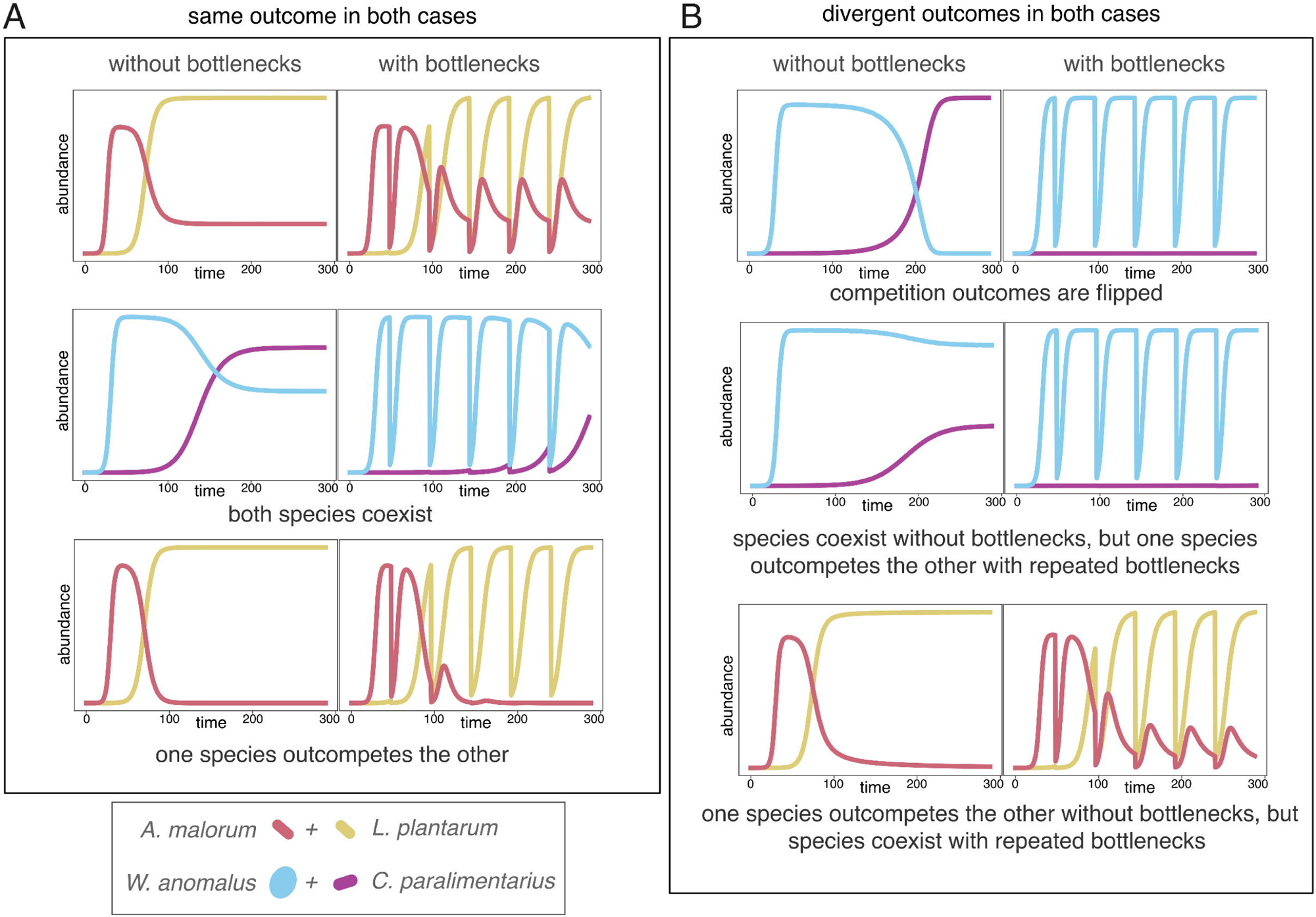
Model predictions of outcomes for selected pairwise competitions between two pairs of species. We compare model growth with and without repeated bottlenecks for the same parameter combinations for the bacterium *C. paralimentarius* and the yeast *W. anomalus*, and the bacteria *A. malorum* and *L. plantarum*. We look at three cases where we observe the same outcome with and without bottlenecks (coexistence and exclusion of one species) and three cases where the addition of bottlenecks changes the outcome of the competition (flipping which species is excluded, changing exclusion to coexistence, or changing coexistence to exclusion).

**Fig 6.**
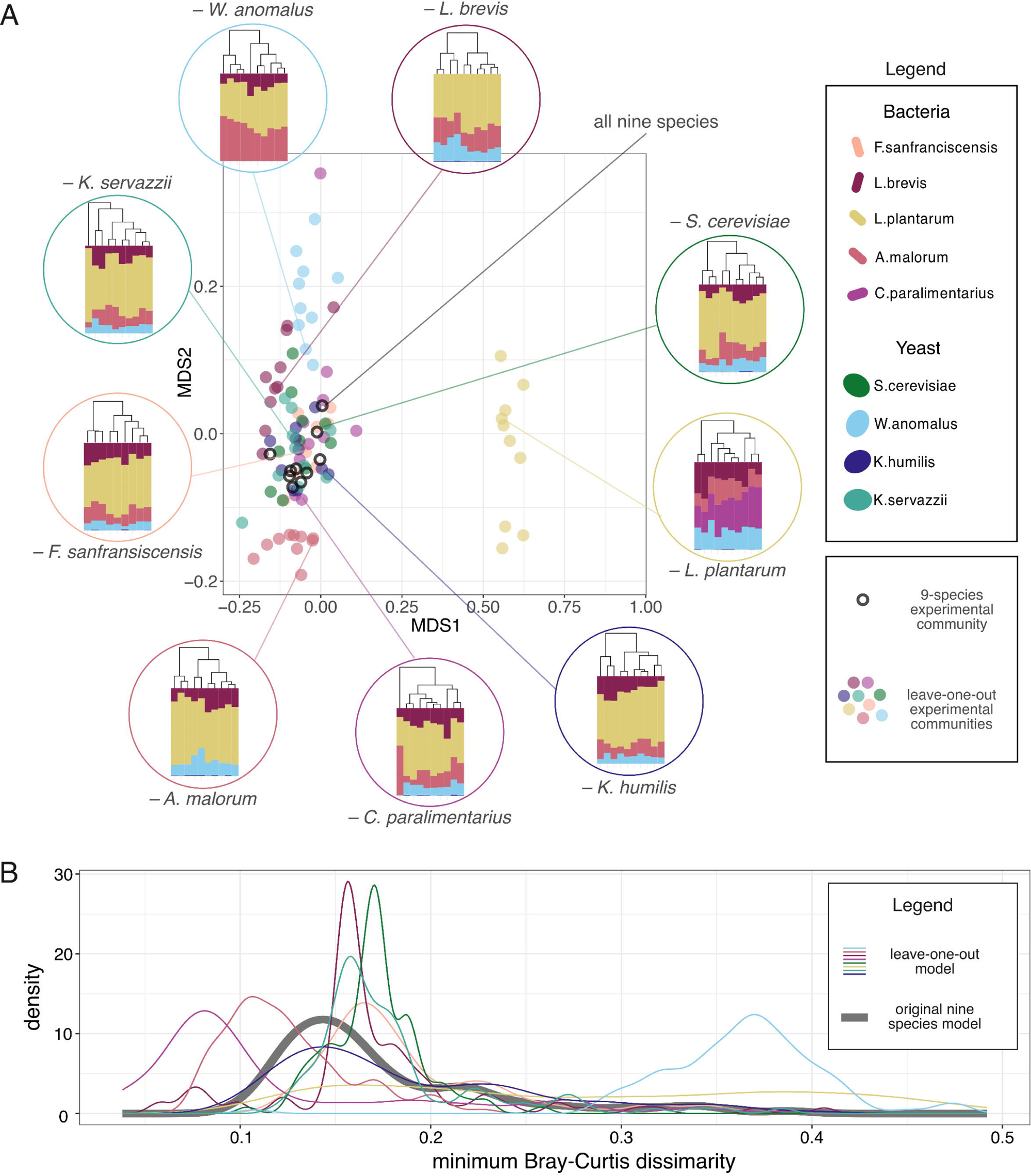
Selected analyses on leave-one-out communities. (A) NMDS plot visualizing Bray-Curtis dissimilarities between all experimentally observed LOO communities. LOO communities are colored by species left out of the community. Inset plots show experimentally observed LOO communities hierarchically clustered by Bray-Curtis dissimilarities. Bar height indicates relative abundance and color indicates species identity. (B) Density plot showing distribution of Bray-Curtis dissimilarities between all model predictions and corresponding experimental communities. The thick gray line indicates the Bray-Curtis dissimilarities between the 9-species model predictions and experimental community. The lines indicating Bray-Curtis dissimilarities between LOO communities and corresponding model predictions are colored by the species left out of the community.

### D. For leave-one-out communities, model predictions and experimental observations align similarly

While our results suggest that the model was able to recapitulate the community outcomes for most species in the pool, we further assessed how the choice of species in the pool might affect composition by performing leave-one-out experiments. We conducted community assembly experiments leaving out one species from the 9-species’ pool and generated corresponding model predictions for each of these communities. We found that community composition remained relatively consistent across most of the leave-one-out communities (LOO) (Fig 4A). With the exception of the community without *L. plantarum*, all LOO communities either had the same species as the 9-species community or lacked only the species that was left out. However, removing *L. plantarum* allowed *C. paralimentarius* to persist in the community. Visualizing community community differences based on Bray-Curtis dissimilarities (Fig 4B), we observed that communities without *L. plantarum* cluster separately on an NMDS plot (p < 0.001, partial R^2^ = 0.81 in PERMANOVA test).

To study whether model communities for the leave-one-outs were more similar to experimental LOOs (as compared to the 9-species model vs experiment), we calculated Bray-Curtis dissimilarities between each model and experimental community for LOOs. We used the gLV model with repeated bottlenecks to obtain predictions for community assembly. Then, we compared the distribution of the minimum Bray-Curtis dissimilarity between one model community and all the corresponding experimental communities (Fig 4C). Most LOO communities had similar distributions of Bray-Curtis dissimilarity, except the ones where *A. malorum* and *C. paralimentarius* were removed, which improved model predictions. However, for communities where *W. anomalus* was removed, model predictions were less accurate in comparison to the predictions of the 9-species community. Overall, the model LOO communities were similar in composition to experimental LOO communities.

## Discussion

Whether the composition of multispecies communities is predictable based on pairwise coexistence or is an emergent property of the community is an ongoing debate amongst microbial ecologists. Studies have provided convincing evidence for both sides of the debate (Friedman, Higgins, and Gore 2017; Venturelli et al. 2018; Dedrick et al. 2023; Momeni, Xie, and Shou 2017; Chang et al. 2023). Our results show that for microbes isolated from sourdough starters, most species’ presence or absence can be reliably predicted based on pairwise interactions. Additionally, in our comparison between model and experiment, we found only two species out of nine for which model predictions differed substantially from experimental observations. This finding suggests that Lotka-Volterra models can be used not only to predict which species can coexist in some microbial communities, but also to predict the abundance of various species in the community.

In addition to our finding that gLV models can provide substantial insight into community assembly in a complex microbial species pool, our work can be understood through the lens of niche-overlap between species pairs. A higher proportion of cross-kingdom species pairs were predicted to coexist than intra-kingdom pairs, a finding that is consistent with empirical patterns observed in prior work on sourdough microbiomes (Landis et al. 2021). Our work shows that niche overlap – based on Chesson’s framework (Letten, Ke, and Fukami 2017) – was lower in bacteria:yeast pairs than yeast:yeast and bacteria:bacteria pairs. Though our experiments do not provide direct insight into the molecular mechanisms that explain pairwise growth outcomes, prior work has demonstrated differences in resource partitioning between bacteria:yeast pairs and sourdough microbes (Minervini et al. 2014; Carbonetto et al. 2018; Oshiro et al. 2021). Inhibition of growth among bacteria:bacteria pairs could be another reason for reduced coexistence (Corsetti, Gobbetti, and Smacchi 1996; Zangeneh, Khorrami, and Khaleghi 2020).

Though there is the potential for substantial complexity underlying species interactions between microbes, our simple nine-species gLV model is able to capture abundance patterns for most species in the pool (Fig 3). Three species (*L. plantarum, L. brevis,* and *W. anomalus*) that were among the four most prevalent in observed communities were also consistently predicted as among the most prevalent in our model communities. Two species that were not observed in the experimental communities (*K. servazzii* and *F. sanfranciscensis*) were consistently predicted to be absent (or extremely low abundance) in model communities, and two species with low abundance in experimental communities (*K. humilis* and *S. cerevisiae*) had consistently low abundance in the models. These outcomes can all be intuited directly from pairwise coexistence outcomes (Fig 2). Species that were present in the final species pool were able to coexist in pairs with other species that were in the final pool. Species that were absent from the species pool were competitively excluded by at least one species that was present in the final pool. We note that this logic breaks down when considering the fate of *F. sanfranciscensis* in leave-one-out communities. Among the species that were observed in the community, only *K. humilis* was predicted to competitively exclude *F. sanfranciscensis*, but removal of *K. humilis* from the pool did not result in the emergence of *F. sanfranciscensis* in models or experiments (Fig 4). The lack of *F. sanfranciscensis* even in the absence of *K. humilis* could represent an emergent effect of the other species on *F. sanfranciscensis* in the multispecies community context – *F. sanfranciscensis* was generally slow-growing and low abundance in most pairs, even though it was often predicted to coexist in pairwise competitions (Fig 2).

Despite the overall success of our pairwise model, there were a few interesting discrepancies between our model and experimental observations. Most strikingly, *C. paralimentarius* was predicted to be present in model communities but was rarely observed, while *A. malorum* was mostly absent from model communities but present in experimental communities. Perhaps due to these discrepancies, model predictions for the LOO communities where these species were removed aligned much better with experimental results (Fig. 6B). Prior studies have often focused on higher-order interactions as potential explanations for discrepancies between pairwise models and experiments. Another possible contributing factor is that most pairwise models do not include demographic processes that are inherent to the growth cycles in both natural and synthetic communities. Microbes are often subjected to periodic bottlenecks in laboratory cultures (Meyer, Kappeli, and Fiechter 1985) as well as in more natural environments like sourdough starters (Van Kerrebroeck, Maes, and De Vuyst 2017; De Vuyst, Comasio, and Van Kerrebroeck 2023). Similar outcomes occur in natural settings due to periodic environmental changes at the daily or seasonal timescale (Williams and Middleton 2008; Shimadzu et al. 2013). Theoretical work suggests that bottlenecks can change coexistence outcomes (Schreiber, Benaïm, and Atchadé 2011; Hening, Nguyen, and Chesson 2021). Since periodic bottlenecks were involved in the community assembly experiment, we decided to investigate whether adding bottlenecks to our model would affect coexistence between microbes in our community. When we added bottlenecks to our model (keeping model parameters constant), we observed that the abundance of *C. paralimentarius* was reduced in competition outcomes (Fig 4, Fig 5), but this did not have a major impact on overall community composition (Appendix S1: Figure S6). One reason for why *C. paralimentarius* performed worse under repeated bottlenecks could be its lower intrinsic growth rate compared to other species, especially *W. anomalus* (Appendix S1: Figure S3). Our results point to demographic processes as a potential explanation for discrepancies between pairwise models and experimental outcomes in some systems. In contrast, demographic processes do not help to explain the absence of *A. malorum* from predicted model communities. In part, this may be due to uncertainty in parameter estimates for *A. malorum*; especially for the *L. plantarum:A. malorum* pair, where the abundance of *A. malorum* declined at later time points and resulted in a wide range of best-fitting growth trajectories. Indeed, one out of one hundred posterior parameter samples resulted in the persistence of *A. malorum* in the bottlenecked community, raising the possibility that small differences in our posterior estimates might have increased its abundance in our model.

Another factor that may explain some of the discrepancies between model predictions and experimental observations could be facilitation of growth between microbes. Constraining parameters to be negative could lead to us ignoring possibilities where the growth of certain species could be facilitated by the presence of other microbes. Mutually stimulating interactions have been documented in other studies, particularly between the yeasts and lactic acid bacteria, or lactic acid bacteria and acetic acid bacteria (Minervini et al. 2014; Sieuwerts, Bron, and Smid 2018; Culp and Goodman 2023). This could potentially explain the persistent lack of *A. malorum* in our model communities, as compared to experimental communities. However, we note that strains do not grow markedly better in co-culture as compared to monoculture growth (Appendix S1: Figure S2). Additionally, *A. malorum* declines in abundance when grown with some of the species in our species pool (notably, *L. plantarum* and *C. paralimentarius*), indicating that its growth is likely not facilitated by the microbes in our system. It is possible that the behaviour of these microbes changes in multi-species communities as compared to pairwise interactions. Better parameter estimates or higher-order interactions might better describe the relationship of *A. malorum* with some species in the community.

Overall, our results suggest that pairwise models are not only sufficient to predict which species will persist in some multi-species communities, but are also able to predict abundances for these species. These results are consistent with prior studies from other systems that have also demonstrated that understanding and manipulating pairwise interactions could be sufficient to understand multi-species’ community dynamics. Several studies in human and mouse gut microbiomes (Marino et al. 2014; Bucci and Xavier 2014; Fisher and Mehta 2014), the cheese microbiome (Mounier et al. 2008) and even sourdough starters (Oshiro et al. 2023) used models derived from the Lotka-Volterra model to identify pairwise inter-species interactions important for community assembly. Additionally, studies have used gLV models or assembly rules based on pairwise dynamics to predict outcomes for community assembly in synthetic communities derived from soil, gut or nasal microbiomes (Friedman, Higgins, and Gore 2017; Venturelli et al. 2018; Dedrick et al. 2023). However, it is unclear why predictions of multi-species communities can be made more accurately in certain systems, while in other systems, mechanism-explicit models or higher-order interactions are used instead (Momeni, Xie, and Shou 2017; Niehaus et al. 2019).

Though conflicting results concerning the utility of pairwise interactions have often been reported (Friedman, Higgins, and Gore 2017; Chang et al. 2023), clearer explanations for disparate results may emerge as more work is undertaken in different microbiomes. For example, heterogeneity in the co-evolutionary histories of species in the pool may drive some differences across studies. Studies indicate that co-evolutionary pressures might be important to maintain coexistence in microbial communities (Hillesland et al. 2014; Barber et al. 2022), and species that co-occur naturally and/or share evolutionary histories might hence be better able to coexist in experimental communities. The widespread prediction of pairwise coexistence between microbes in our species pool may be the reason that the multi-species gLV model predicted species’ presence and abundance. Additional studies or meta-analyses of studies applying gLV model predictions to experimental results could help clarify the underlying causes of these discrepant results and provide substantial insights into the processes governing community assembly in nature.

## Acknowledgements

We would like to thank Dr. Angela Oliverio, Dr. Daniel Wuitchik, Nicolas Louw, Alejandro Calderón and members of the Wolfe lab for their helpful comments on the manuscript.

