## Appendix S1 for "Pairwise interactions and serial bottlenecks help explain species composition in a multi-species microbial community"

**Section S1. Derivation of formulae for $\rho$ and $\frac{f_2}{f_1}$ :**

The definitions of niche overlap and fitness difference as given by Chesson are -

$$\rho = \sqrt{\frac{\alpha_{12} * \alpha_{21}}{\alpha_{11} * \alpha_{22}}}$$
$$\frac{f_2}{f_1} = \sqrt{\frac{\alpha_{11} * \alpha_{12}}{\alpha_{21} * \alpha_{22}}} \tag{1}$$

The generalized Lotka-Volterra (gLV) equations used by Chesson have the following form -

$$\frac{dN_i}{N_i dt} = r_i(1 - \alpha_{ii}N_i - \sum_{j \neq i} \alpha_{ij}N_j) \tag{2}$$

Whereas, the gLV equations we use in this paper have the following form -

$$\frac{dN_i}{N_i dt} = r_i + a_{ii}N_i + \sum_{j \neq i} a_{ij}N_j \tag{3}$$

Comparing equations 2 and 3, we get the following equivalences between $\alpha_{ii}$ and $a_{ii}$, and $\alpha_{ij}$ and $a_{ij}$ -

$$\alpha_{ii} = -a_{ii} * r_i$$
$$\alpha_{ij} = -a_{ij} * r_i \tag{4}$$

Replacing all instances of $\alpha_{11}$, $\alpha_{12}$, $\alpha_{21}$,$\alpha_{22}$ by $a_{11}$, $a_{12}$, $a_{21}$, $a_{22}$ in the formulae in 1, we get the following formulae for niche overlap and fitness difference -

$$\rho = \sqrt{\frac{\alpha_{12} * \alpha_{21}}{\alpha_{11} * \alpha_{22}}}$$
$$\frac{f_2}{f_1} = \sqrt{\frac{\alpha_{11} * \alpha_{12}}{\alpha_{21} * \alpha_{22}}} * \frac{r_2}{r_1} \tag{5}$$

These formulae were used in the calculations of niche overlap and fitness difference.

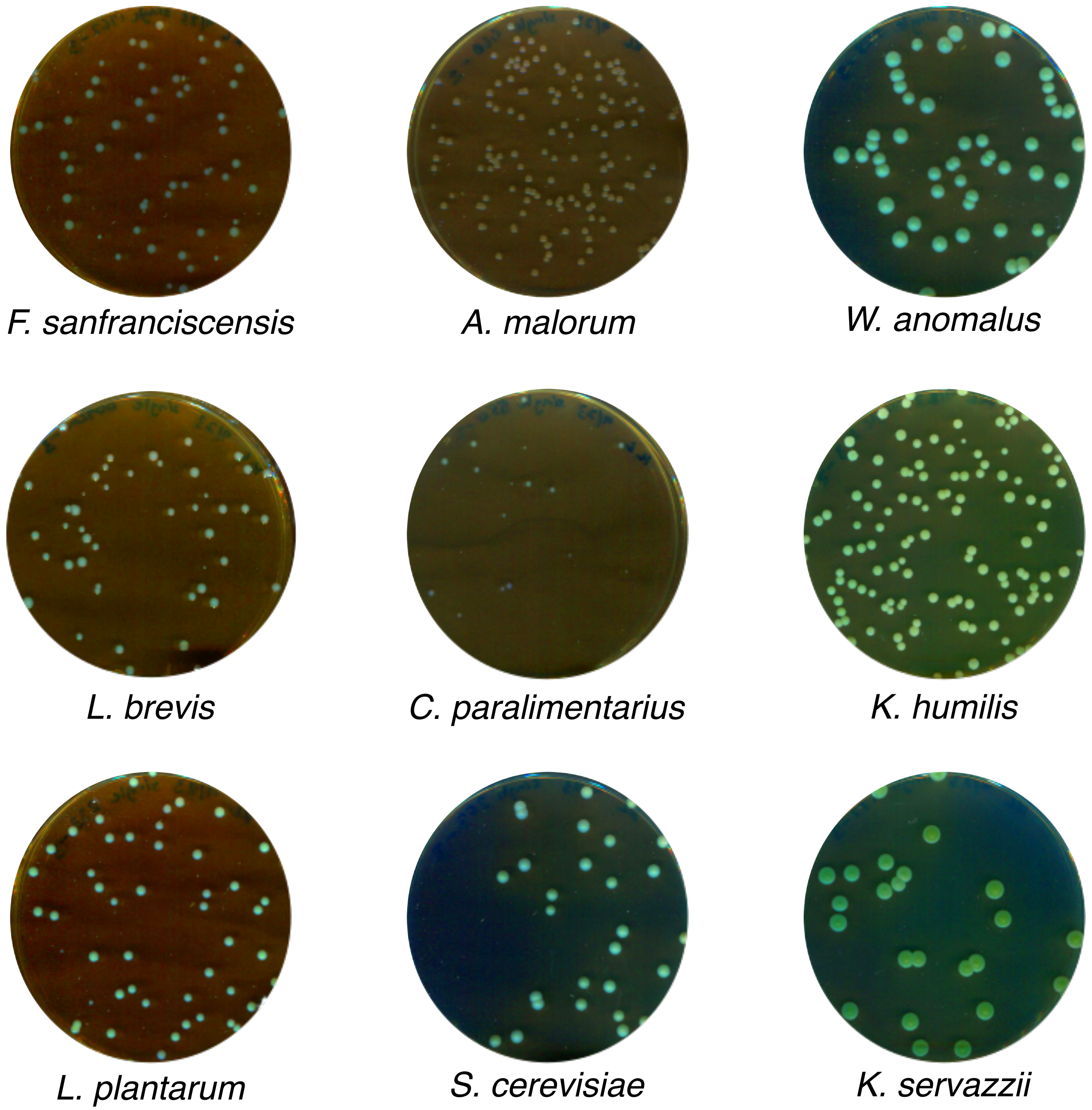


**Figure S1. Examples of bacterial and yeast colonies.** These images indicate differences in colony morphologies by which we differentiated between species in our community.

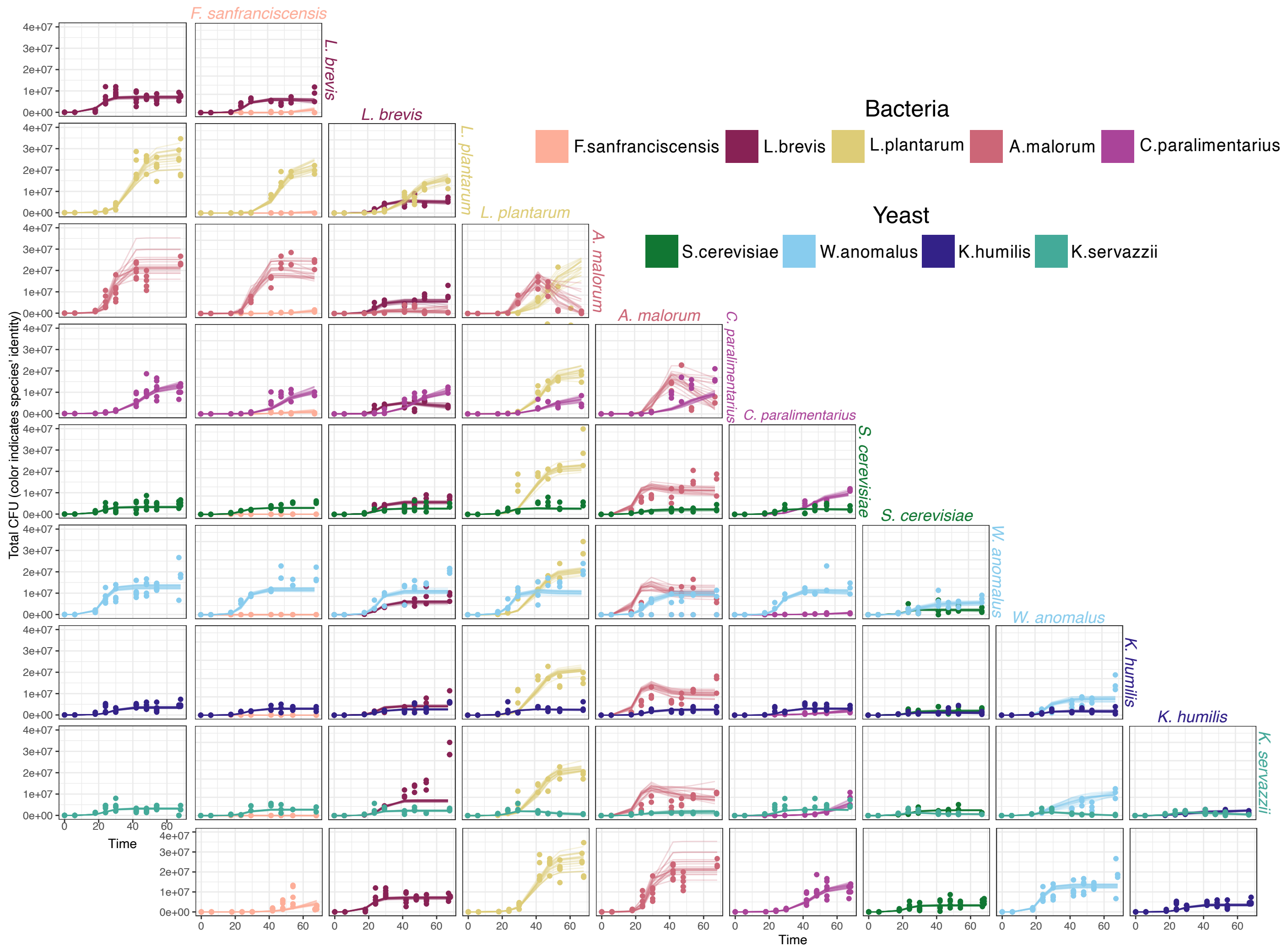


**Figure S2. Measured and extracted growth curves for all species by themselves and for all pairwise combinations of species.** Single-species growth curves for corresponding species are depicted in the first column on the left and the first row from the bottom. Points indicate experimental observations of CFU counts, lines indicate best-fitting growth curves which were used for calculating growth rates and interaction coefficients.

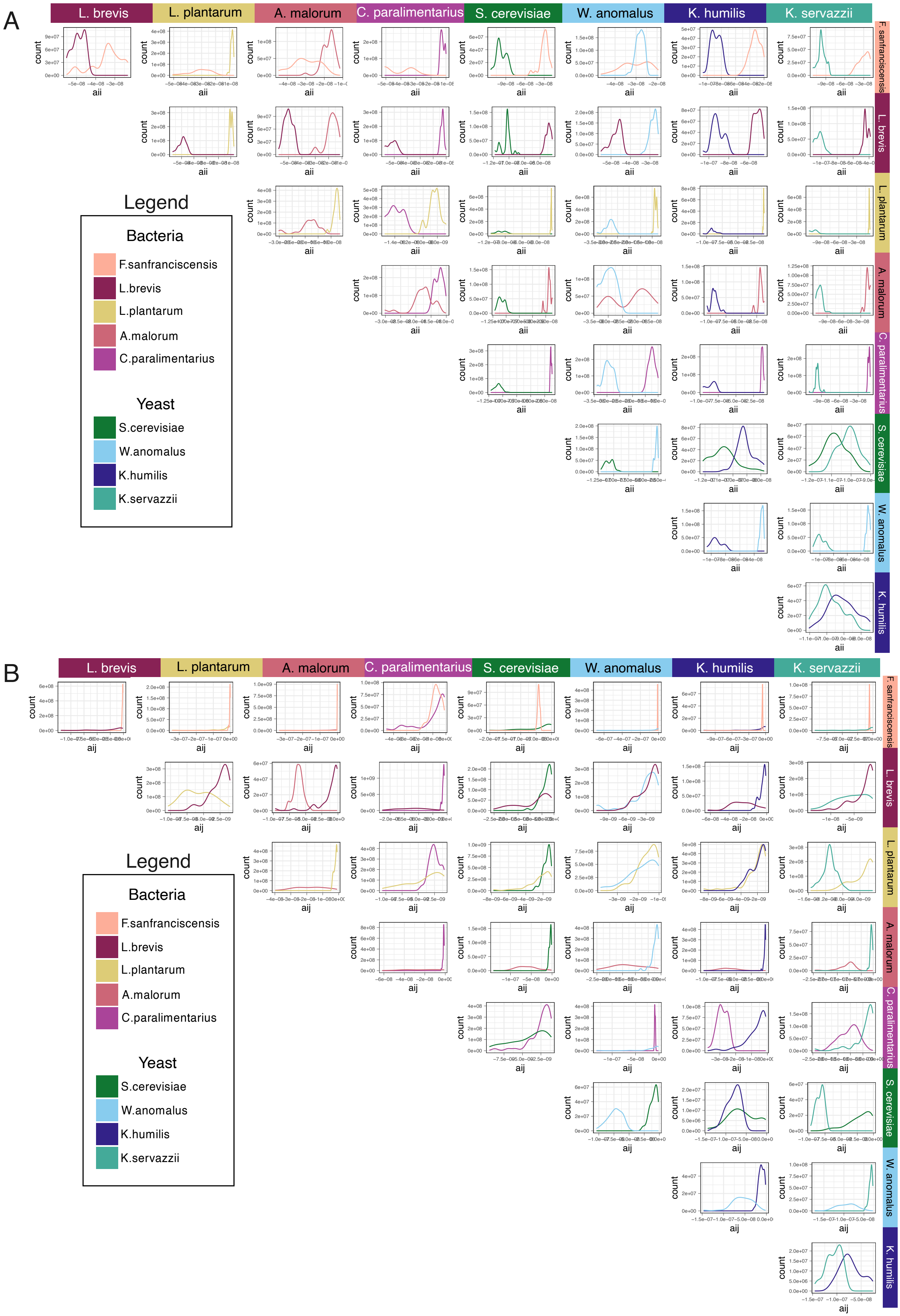
A
L. brevis
L. plantarum
A. malorum
C. paralimentarius
S. cerevisiae
W. anomalus
K. humilis
K. servazzii
F. sanfranciscensis
count
aii
Legend
Bacteria
F.sanfranciscensis
L.brevis
L.plantarum
A.malorum
C.paralimentarius
Yeast
S.cerevisiae
W.anomalus
K.humilis
K.servazzii
B
aij

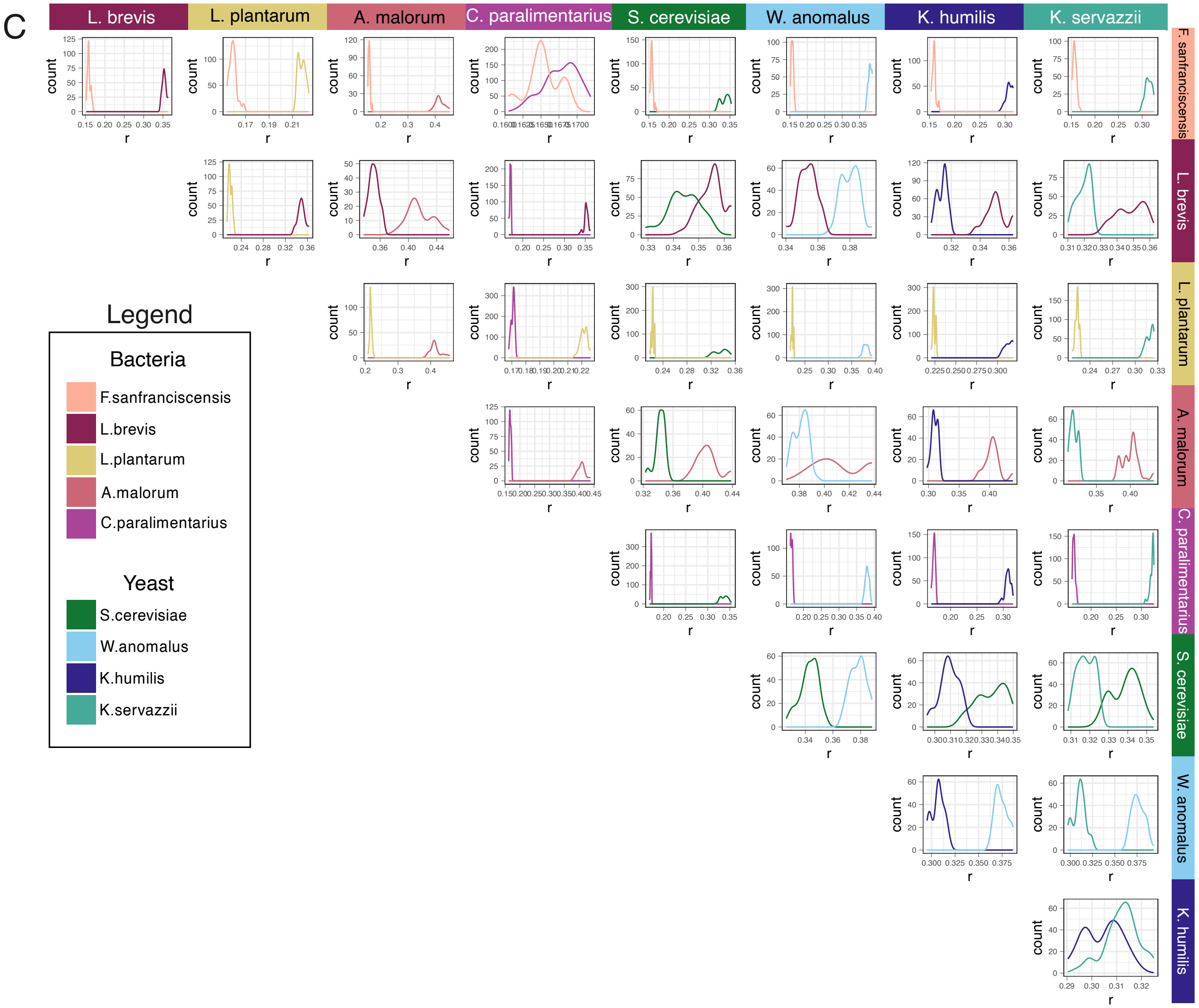


**Figure S3. Distributions of the growth parameters $r_i$, $a_{ii}$, and $a_{ij}$.** Distributions of values of (A) $a_{ii}$, (B) $a_{ij}$, and (C) $r_i$ sampled from experimentally observed growth curves (Figure S2). These parameter values were used for all model predictions. For all figures, colors indicate species for which the given parameter was measured. Parameter values are on the x-axis, and a representation of number of parameters having that value on the y-axis.

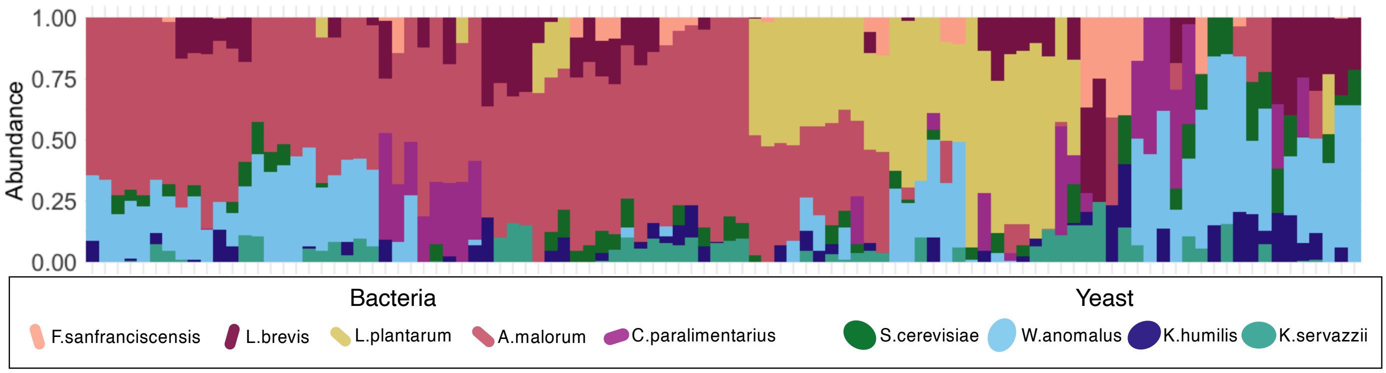


**Figure S4. Model predictions with randomized pairwise interactions.** Model communities hierarchically clustered by Bray-Curtis dissimilarities. Height of each bar indicates relative abundance of corresponding species in a given community.

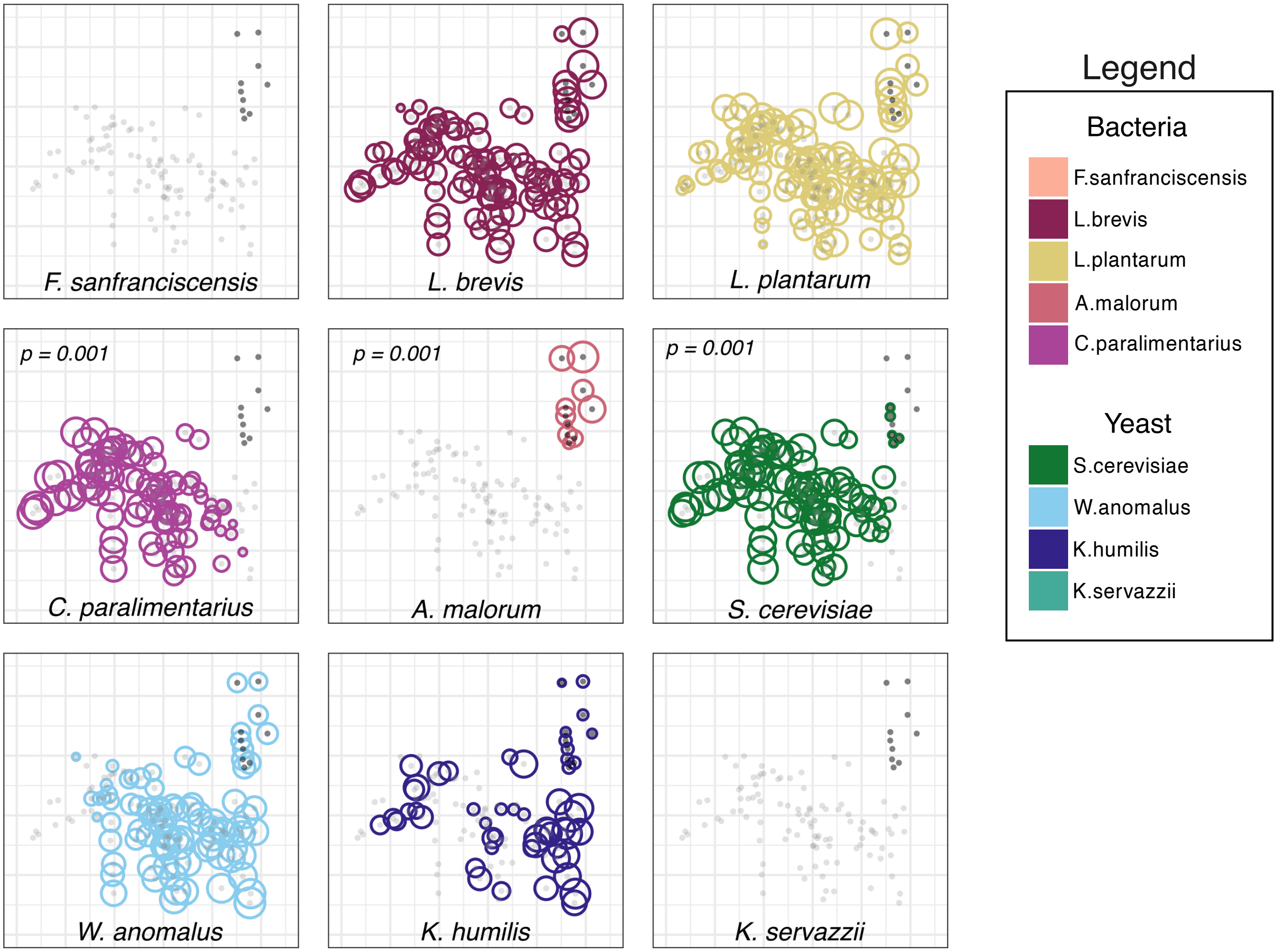


**Figure S5. Indicator species analysis for all species in the 9-species community.** Each graph shows the abundance of one species overlaid on top of the NMDS plot showing the distribution of model and experimental communities. The middle row shows the three species which were indicators for either model or experimental communities. Light gray dots indicate model communities, dark gray dots indicate experimental communities. Size of colored circles indicates abundance of species in the corresponding community.

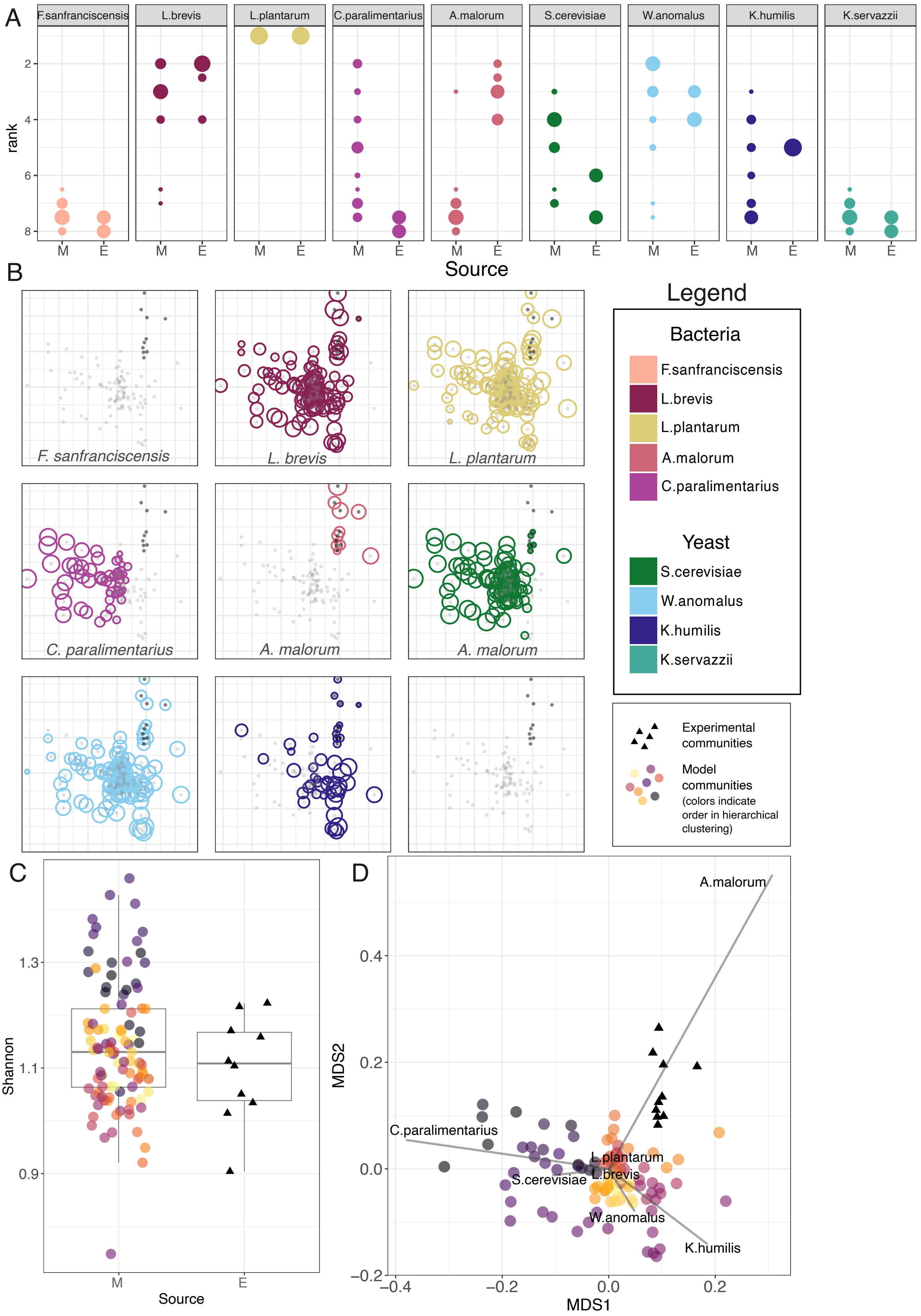
A
F.sanfranciscensis
L.brevis
L.plantarum
C.paralimentarius
A.malorum
S.cerevisiae
W.anomalus
K.humilis
K.servazzii
rank
2
4
6
8
M
E
Source
B
F. sanfranciscensis
L. brevis
L. plantarum
C. paralimentarius
A. malorum
A. malorum
Legend
Bacteria
F.sanfranciscensis
L.brevis
L.plantarum
A.malorum
C.paralimentarius
Yeast
S.cerevisiae
W.anomalus
K.humilis
K.servazzii
Experimental communities
Model communities (colors indicate order in hierarchical clustering)
C
Shannon
1.3
1.1
0.9
M
E
Source
D
A.malorum
C.paralimentarius
L.plantarum
S.cerevisiae
L.brevis
W.anomalus
K.humilis
MDS2
0.4
0.2
0.0
−0.2
−0.4
−0.2
0.0
0.2
MDS1

**Figure S6. Selected analyses for model predictions with serial bottlenecks.** (A) Distributions of species' ranks compared between model (M) and experimental (E) communities. Higher ranks indicate greater abundance in the community. Dot size indicates the fraction of communities where species had corresponding ranks.(B) Indicator species analysis for all species in the community. Each graph shows the abundance of one species overlaid on top of the NMDS plot (D) showing the distribution of model and experimental communities. The middle row shows the three species which were indicators for either model or experimental communities. Light gray dots indicate model communities, dark gray dots indicate experimental communities. Size of colored circles indicates abundance of species in the corresponding community. (C) Shannon diversity index between M and E communities. (D) NMDS plot visualizing Bray-Curtis dissimilarities between M and E communities. For D and E, black triangles indicate experimental communities and coloured circles indicate model communities. Colors indicate distance in hierarchical clustering from Fig 3A for model communities. Gray vectors and species names indicate species' scores.
